# The Hunt for Cholesteryl Ester Hydrolases: Identification of Lipoprotein Lipase as a Cholesterylesterase

**DOI:** 10.64898/2026.07.02.736233

**Authors:** Aakash Chandramouli, Siddhesh S. Kamat

**Author notes:** To whom the correspondence should be made.

## Abstract

Cholesteryl esters (CEs) are central intermediates in cholesterol storage and transport, yet the enzymes responsible for their hydrolysis in mammals remain poorly defined. While lysosomal acid lipase is the only well-established acidic CE hydrolase, the molecular identity of physiologically relevant neutral CE hydrolases has remained unresolved. Here, we systematically profiled CE hydrolase activity across mouse tissues and blood using substrate-based LC-MS assays, tissue fractionation, and inhibitor screening. We observed robust CE hydrolase activity in multiple tissues and circulation, with activity predominantly enriched in membrane fractions and strongly sensitive to broad-spectrum metabolic serine hydrolase inhibitors. Pharmacological screening excluded previously proposed neutral CE hydrolases, including NCEH1 and LIPE, and identified tetrahydrolipstatin-sensitive lipoprotein lipase (LPL) as a candidate CE hydrolase. Competitive activity-based protein profiling analyses in RAW264.7 macrophages further supported selective enrichment and inhibition of LPL. Biochemical characterization demonstrated that recombinant wild-type LPL, but not the catalytic S159A variant, efficiently hydrolyzed CEs *in vitro*. Importantly, this activity required co-expression of the lipase maturation factor 1, indicating that LPL-mediated CE hydrolysis is dependent on proper enzymatic maturation. Together, these findings identify LPL as a previously unrecognized mammalian CE hydrolase and expand its functional role beyond triglyceride metabolism.

## INTRODUCTION

Cholesterol is a ubiquitous and essential lipid present in all animal tissues, where it serves both structural and metabolic roles (1). Structurally, cholesterol is a 27-carbon sterol composed of a rigid tetracyclic steroid backbone, a hydroxyl group at the C3 position, and an iso-octyl side chain at C17 (**Figure 1**) (2). In biological systems, cholesterol exists in two principal forms: unesterified (or “free”) cholesterol and cholesteryl esters (CEs), in which the C3 hydroxyl group is esterified with fatty acids of varying lipid tails. This interconversion between free and esterified cholesterol is central to cellular lipid homeostasis and is tightly regulated by enzymatic processes (**Figure 1**) (2).

**Figure 1.**
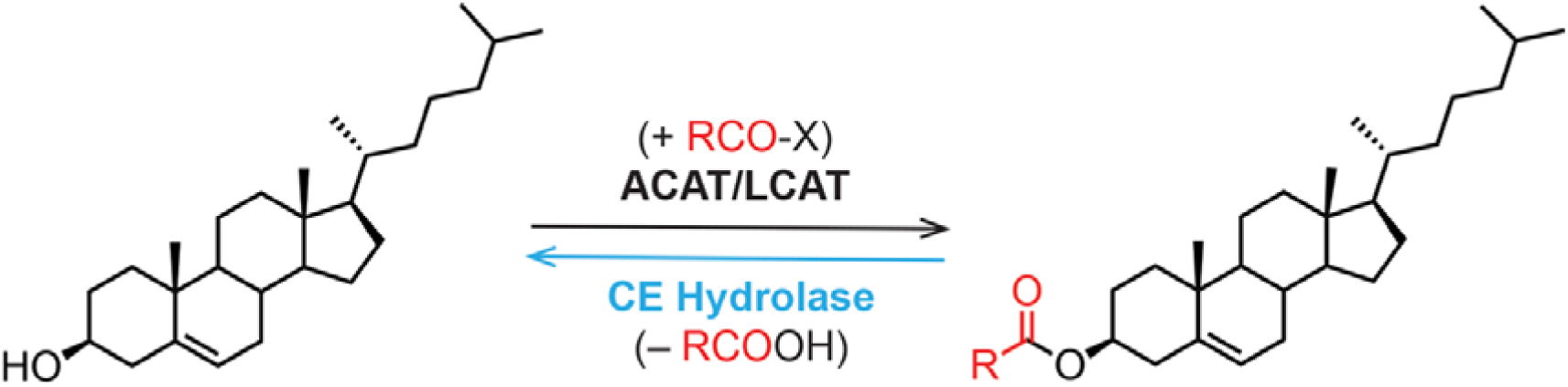
Metabolism of cholesteryl esters (CEs). ACAT and LCAT biosynthesize CEs from free cholesterol using acyl-CoA (for ACAT) or phosphatidylcholine (for LCAT) as the source of the fatty acids. Counter to this, a CE hydrolase makes free cholesterol from CEs via a hydrolytic reaction.

Cholesterol esterification is catalyzed primarily by two enzymes: acyl-CoA:cholesterol acyltransferase (ACAT) and lecithin:cholesterol acyltransferase (LCAT). ACAT operates within cells, utilizing fatty acyl-CoA substrates to generate CEs that are subsequently stored in lipid droplets (3, 4). In contrast, LCAT functions in the extracellular environments, particularly in plasma, where it transfers fatty acyl chains, predominantly from the *sn-2* position of abundant phospholipids like phosphatidylcholine (PC), to cholesterol (5). Through these complementary activities, ACAT and LCAT maintain the dynamic balance between free cholesterol and its esterified form across cellular and systemic compartments.

Functionally, cholesterol is indispensable for maintaining membrane integrity and fluidity. Its ability to modulate lipid packing contributes to the formation of specialized membrane microdomains, often referred to as lipid rafts, which facilitate intracellular trafficking and signal transduction (1, 6). Beyond its structural role, cholesterol also serves as a precursor for synthesizing bile acids, steroid hormones, and vitamin D, underscoring its metabolic versatility (6). Given these diverse functions, intracellular cholesterol levels are under stringent control. Excess free cholesterol, which can be cytotoxic, is rapidly esterified and stored as CEs within lipid droplets. In contrast to the well-established roles of free cholesterol, cholesteryl esters have traditionally been regarded as relatively inert storage forms. They function as reservoirs of both cholesterol and fatty acids and play a key role in lipid transport through lipoprotein particles such as high-density lipoproteins (HDL) and low-density lipoproteins (LDL). However, this view has evolved significantly in recent years. Accumulating evidence implicates CE accumulation in the pathogenesis of several diseases, including atherosclerosis, neurodegenerative disorders, and cancer (7–11). Moreover, CEs are susceptible to oxidative modification, giving rise to oxidized CEs, which can interact with proteins and influence diverse signaling pathways (12–15). These findings highlight that CEs are not merely passive storage molecules but may actively participate in cellular regulation and disease progression.

While the enzymology of cholesterol esterification has been extensively characterized, comparatively little is known about the enzymes responsible for CE hydrolysis (**Figure 1**). This process, which regenerates free cholesterol and fatty acids from stored esters, is essential for maintaining lipid flux and cellular homeostasis. Over the past several decades, multiple candidate enzymes have been proposed as CE hydrolases in mammals, broadly classified based on their pH optima into acidic and neutral CE hydrolases. Among acidic enzymes, lysosomal acid lipase (LAL) remains the only well-established CE hydrolase in mammals (16, 17). Localized to lysosomes, LAL catalyzes the breakdown of both CEs and triacylglycerols delivered via endocytosed lipoproteins. Genetic deficiency of LAL leads to severe lipid storage disorders, including Wolman disease and CE storage disease (CESD), both characterized by the pathological accumulation of CEs in multiple tissues (16, 17). These observations underscore the critical role of LAL in intracellular lipid catabolism.

In contrast, the identity and physiological relevance of neutral CE hydrolases have remained ambiguous (18). Two enzymes, hormone-sensitive lipase (HSL, encoded by LIPE) and neutral cholesteryl ester hydrolase 1 (NCEH1, also known as KIAA1363 or AADACL1), have been studied in this context (19–22). HSL was initially characterized as a triacylglycerol lipase but was later shown to exhibit CE hydrolase activity *in vitro* and in certain tissues. However, genetic ablation of LIPE in mice does not result in alterations in CE levels, raising questions about its role as a physiologically relevant CE hydrolase. Similarly, NCEH1 was first identified as an enzyme that hydrolyzes 2-acetyl monoalkylglycerol ether in cancer cells and was subsequently reported to act on a broader range of lipid esters, including CEs. Yet, NCEH1 knockout animal and cellular models also fail to show substantial changes in CE hydrolysis or accumulation. Collectively, these findings suggest that neither HSL nor NCEH1 alone accounts for the bulk of neutral CE hydrolytic activity in mammals, implying the existence of additional, as of yet unidentified, enzymes.

To address this gap, we systematically investigated the presence and molecular identity of CE hydrolase activity across mammalian tissues and blood. Previous studies have largely focused on adipose tissue and macrophages, despite the widespread distribution of CEs in diverse tissues. Using a combination of tissue fractionation, substrate-based hydrolysis assays, and pharmacological inhibition, we demonstrate that CE hydrolase activity is broadly distributed across different mouse tissues, with the notable exception of skeletal muscle, and is also detectable in circulation. Biochemical profiling further indicates that this activity is mediated by a member of the metabolic serine hydrolase enzyme family. To identify the enzyme responsible, we employed inhibitor-based screening coupled with competitive chemoproteomic approaches. These analyses converged on lipoprotein lipase (LPL) as a candidate CE hydrolase. LPL is classically known for its role in hydrolyzing triacylglycerols in circulating lipoproteins, thereby facilitating fatty acid uptake into tissues. Our findings expand this functional repertoire by implicating LPL in CE metabolism. Biochemical characterization of recombinant LPL confirms its ability to hydrolyze CEs. Importantly, we find that this activity in cells requires the presence of lipase maturation factor 1 (LMF1), a chaperone essential for the activation of several lipases, including LPL.

Together, these results support a model in which LPL, in concert with LMF1, contributes to neutral CE hydrolysis in mammalian systems. This functional annotation provides a mechanistic framework for understanding how CE pools are mobilized across tissues and suggests a broader role for LPL in lipid homeostasis beyond its established function in triglyceride metabolism. By identifying a previously unappreciated pathway for CE hydrolysis, this work advances our understanding of cholesterol metabolism and opens new avenues for exploring the role of lipid dynamics in physiology and disease.

## MATERIALS AND METHODS

### Materials

Unless otherwise mentioned, all chemicals, buffers, and reagents were purchased from Sigma-Aldrich (Merck). All tissue culture media and supplies were purchased from HiMedia. All solvents used for mass spectrometry were LC-MS grade and were purchased from JT Baker (Acetonitrile) or ThermoFisher Scientific (Methanol and Isopropanol).

### Mammalian cell culture and treatments

All mammalian cell lines used in this study were purchased from ATCC and cultured in Dulbecco’s Modified Eagle’s Medium (DMEM) containing 10% (v/v) Fetal Bovine Serum (FBS) and 1% (v/v) Penicillin-Streptomycin (MP Biomedicals) at 37 °C and 5% (v/v) CO_2_. All cell lines described in this study were routinely stained with 4’,6-diamidino-2-phenylindole and visualized by microscopy to ensure that they were devoid of any mycoplasma contamination (23, 24). For lipid measurements (per biological replicate), the cells were treated with the inhibitor or vehicle (DMSO) at respective concentrations and treatment time (specified in corresponding figure legends) in a single 10 cm tissue culture dish, with the cells at 80 – 85% confluence. Post-treatment, the cells were washed with cold sterile Dulbecco’s phosphate-buffered saline (DPBS) (3x), harvested by scraping, and centrifuged at 300g for 5 min at 4 °C to get a cell pellet. The harvested cell pellets were flash-frozen using liquid nitrogen and stored at −80 °C until further use.

### Tissue and blood samples from mice

All protocols involving mice used for this study were approved by the Institutional Animal Ethics Committee at IISER Pune as per guidelines provided by the Committee for the Purpose of Control and Supervision of Experiments on Animals (CPCSEA) constituted by the Government of India (No: IISER_Pune IAEC/2023_03/02). All mice used in this study were from the C57Bl/6J strain and between 10 – 12 weeks of age. Briefly, the mice were deeply anesthetized, and blood samples (∼ 500 μL) were collected via the retro-orbital route into tubes containing 4% (w/v) EDTA solution (for whole blood and plasma) or in tubes without EDTA (for serum). Thereafter, whole blood samples were immediately flash-frozen, or were further processed for serum and plasma separation as per established protocols and subsequently flash-frozen using liquid nitrogen and stored at −80 °C until further use. Following this, mice were euthanized by cervical dislocation, and the various organs were harvested. All organs were washed with cold sterile DPBS (2x), transferred to 1.5 mL microcentrifuge tubes, weighed, and flash-frozen using liquid nitrogen and stored at −80 °C until further use.

### Preparation of tissue fractions

Tissue fractions were prepared as per established protocols as reported earlier (25). The pre-weighed tissues were thawed on ice, resuspended in 500 μL of cold sterile DPBS, and homogenized using a Bullet Blender 24 (Next Advance) using one scoop of 0.5-mm diameter glass beads (Next Advance) for brain, or one scoop of 2.3-mm diameter stainless steel beads (Next Advance) for the liver and heart and one scoop of 1-mm diameter zirconium oxide beads (Next Advance) for the kidneys, spleen, and lungs, at a speed setting of 8 for 3 min at 4 °C (2x). To this tissue homogenate, 500 μL of cold sterile DPBS was added and mixed by vortexing. These homogenates were subsequently probe sonicated for 90 sec (2 sec ON and 3 sec OFF pulses at 60% amplitude) using a medium-sized probe. Thereafter these homogenates were centrifuged at 1000g for 5 min at 4 °C to separate the beads and tissue debris from the tissue proteome. A small part of the resulting tissue lysate (supernatant, ∼100 µL) was separated by pipetting and labeled as tissue lysate and the remaining tissue lysate (supernatant, ∼700 μL) was separated by pipetting and centrifuged at 100,000g for 1 h at 4 °C in an Optima MAX-XP ultracentrifuge (Beckman Coulter). Following ultracentrifugation, the resulting supernatant (∼400 μL) was separated by pipetting and labeled as the soluble fraction for a particular tissue. The resulting pellet was washed with cold sterile DPBS (3x) and resuspended in 400 μL cold sterile DPBS by sonication. This resulting lysate was labeled as the membrane fraction for a particular tissue. The protein concentrations of the tissue lysate, and the membrane and soluble proteomic fractions were measured using the Pierce BCA Protein Assay Kits (ThermoFisher Scientific, catalog no.: 23225).

### CE hydrolase substrate assays

For the CE hydrolase substrate assays, 40 µg of lysates (tissue or cells) were incubated with 100 µM of C18:1 CE (Sigma, catalog no.: C9253) substrate presented in the form of vesicles with phosphatidylcholine [C18:0/18:0 PC (DSPC), Avanti Polar Lipids, catalog no.: 850365C] mixed in a ratio of 1:4 (mol/mol) (CE:PC), and 100 µM of sodium taurocholate in DPBS to a final volume of 100 µL at 37 °C with constant shaking for 1 h. For all assays with blood (whole, plasma or serum), 20 µL of the sample was taken and reaction was adjusted to a final volume of 100 µL. Post 1 h incubation, the reaction was quenched with the addition of 300 µL of 2:1 (v/v) chloroform:methanol (CHCl_3_:MeOH) containing 0.5 nmol of internal standard [C15:0 free fatty acid (FFA), Sigma, catalog no.:P6125]. The organic and the aqueous phases were separated by centrifugation at 1500g for 15 min. The organic phase (bottom) was collected in a new glass vial, dried under a stream of N_2_ gas, and resolubilized with 300 µL of CHCl_3_. Next, 80 µL of this mixture was mixed with 40 µL of MeOH, and 5 µL of this was used for LC-MS analysis.

The LC-MS detection and quantification of the free-fatty acid product was carried out using Single Ion Monitoring (SIM) mode on an Agilent G6125B single quadrupole LC/MS coupled with an Agilent 1260 Infinity II UHPLC system (25). Reverse-phase liquid chromatography (LC) separation was achieved using a Luna 5U C5 column (Phenomenex, 5 μm, 50 x 4.6 mm, catalog no.: 00B-4043-E0) coupled to guard column (Phenomenex, 3.2 x 8 mm, catalog no.: KJ0-4282). The flow rate for the LC method was 0.5 mL/min, with the composition of solvent A being 95:5 (v/v) H_2_O:MeOH, and that of solvent B being 60:35:5 (v/v) isopropanol:MeOH:H_2_O. A typical LC-run was 21 min, with the following gradients for elution: 0 – 1.5 min, 10% solvent B; 1.5 – 5 min, 10 – 100% linear gradient of solvent B; 5 – 18 min, 100% solvent B (wash step); 18 – 18.1 min, 100 – 10% solvent B; 18.1 – 21 min, 10% solvent B (re-equilibration step). The temperature of the autosampler (samples) and column were maintained 8 °C and 40 °C respectively. All LC-MS acquisitions were performed using an electrospray ionization source in the negative ion mode, with the following MS parameters: drying gas temperature at 250 °C, drying gas flow at 10 L/min, nebulizer pressure at 45 psi, quad temperature at 100 °C, capillary voltage at 4 kV, and fragmentor voltage at 75 V. SIM ions for product C18:1 FFA (m/z 281.2) and internal standard C15:0 FFA (m/z 241.2) were monitored for quantification. The product release was quantified by measuring the area under the curve for the peak corresponding to C18:1 FFA (produced from C18:1 CE) and normalizing it to the internal standard (C15:0 FFA). The non-enzymatic substrate hydrolysis rate was obtained using heat-denatured protein (15 min at 95°C, followed by cooling at 4°C for 10 min, 3x) from the respective origin as a comparison control as reported earlier.

### Targeted measurement for cholesterol and CEs

Cholesterol and CEs were extracted from vehicle or inhibitor treated cells using a modified Folch extraction procedure reported by us earlier (2). Dried lipid extracts were dissolved in 200 µL CHCl_3_:MeOH (2:1, v/v), and 10 µL aliquots were injected for LC–MS/MS analysis. LC-MS analyses were performed on an Agilent 6470 Triple Quadrupole LC/MS using established multiple reaction monitoring transition-based analysis for cholesterol and CEs previously reported by us (2). All LC and MS instrument parameters and acquisition settings were identical to previously validated laboratory protocols to ensure reproducibility and sensitivity (2). Cholesterol and CEs were quantified by normalizing their respective areas under the curve and normalizing it to the area under the curve of the internal standard [CE17:0 (Avanti Polar Lipids, catalog no.: 700186M) or cholesterol-d7 (Avanti Polar Lipids, catalog no.: 700041P)] and then normalizing to total cellular proteins.

### Competitive chemoproteomics

After treatment with inhibitors, the cells were harvested by scraping and lysed by sonication as reported earlier (23). In this experiment, whole cell lysates were used (without fractionation). For proteomic sample preparations, cell lysates (2Cmg/mL in 1CmL DPBS) were labeled with FP-biotin (10 μM, 1 h, 37 °C with shaking). After labeling, proteins were denatured and reductively alkylated using iodoacetamide as previously reported. Proteolytic digestion was carried out overnight at 37 °C using sequencing-grade trypsin (Promega, catalog no.: V5111) at an enzyme-to-substrate ratio of 1:50 (w/w). Tryptic peptides were tagged by reductive dimethylation as per protocols described earlier with light (H_2_CO) (Sigma-Aldrich; catalog no.: 252549) or heavy (D_2_CO) formaldehyde (Cambridge Isotope Laboratories Inc.; catalog no.: DLM-805-20) and sodium cyanoborohydride (NaBH_3_CN) (23, 26). Tryptic peptides from the inhibitor-treated group (light labeled) and DMSO-treated group (heavy labeled) were mixed, desalted with the StageTip protocol (27) and dried under vacuum prior to LC–MS/MS analysis.

All proteomic LC–MS/MS analyses were conducted on a Sciex TripleTOF 6600 mass spectrometer coupled to an Eksigent nanoLC 425 system (26, 28). Peptides were initially loaded onto a C18 trap column and subsequently separated on a C18 analytical column (15 cm x 75 µm internal diameter) using a linear acetonitrile gradient at a constant flow rate of 300 nL/min. Solvent A consisted of water with 0.1% formic acid, and solvent B consisted of acetonitrile with 0.1% formic acid. A typical gradient program included 5% solvent B for 1 min, a linear increase to 30% solvent B over 330 min, 90% solvent B for 20 min, followed by re-equilibration at 5% solvent B for 10 min. Data were acquired in information-dependent acquisition mode across an m/z range of 200–2,000. Each full MS scan was followed by MS/MS acquisition of the 15 most intense precursor ions. Dynamic exclusion parameters were set to a repeat count of 2 and an exclusion duration of 6 sec. Peptide identification was performed using ProteinPilot (v2.0.1, Sciex) with the Paragon and ProGroup algorithms against a *Mus musculus* RefSeq protein database (Release 109).

Carbamidomethylation of cysteine was set as a fixed modification, while methionine oxidation and N-terminal acetylation were considered variable modifications. Precursor and fragment mass tolerances were set to 20 ppm and 50 ppm, respectively. A decoy database strategy was employed to control false discovery rates (FDR), and only identifications with FDR < 1% were retained. For reductive-dimethylation-based quantification of tryptic peptides, the N-terminal and lysine dimethyl labels were specified as fixed modifications in the search algorithm within ProteinPilot software for quantification of identified proteins and the enrichment ratio (heavy label to light label) was used for filtering serine hydrolase (SH) enzymes for subsequent analysis. For a SH enzyme to be considered for further analysis, it needed to be identified in at least two biological replicates and have one or more quantified peptides per replicate it was identified in.

### Gel-based ABPP assays

All gel-based ABPP assays were performed using protocols previously reported by us (25). Briefly, 100 μL of 2 mg/mL lysates from mammalian cells were incubated with 2 μM FP-rhodamine for 45 min at 37 °C with constant shaking (Eppendorf, ThermoMixer C). The reactions were then quenched by adding 34 μL of 4x SDS-PAGE loading buffer, followed by boiling for 10 min. Finally, 30 μL of this sample was loaded on a 12.5% SDS-PAGE gel. Competitive gel-based ABPP experiments were done as reported earlier. All gels were visualized using an iBright1500 gel imaging system (Invitrogen).

### Cloning and overexpression of LPL and LMF1 in HEK293T cells

Total RNA was extracted from RAW264.7 cells using TRIzol reagent (Invitrogen, catalog no.: 15596018) as per manufacturer’s instructions. Fresh cDNA was prepared from total RNA using high-capacity cDNA reverse transcription kit (Applied Biosystems, catalog no.: 4374966) as per manufacturer’s protocol. LPL and LMF1 were amplified from cDNA and cloned into pcDNA3.1 myc-HisA(-) vector between XbaI and KpnI restriction sites (for LPL), and between XhoI and HindIII restriction sites (for LMF1), respectively. The S159A variant of LPL was generated by site-directed mutagenesis using Phusion Plus DNA polymerase (Invitrogen, catalog no.: F630S) and DpnI (New England Biolabs, catalog no.: R0176L) as per manufacturer instructions.

HEK293T cells were cultured in RPMI1640 medium supplemented with 10% (v/v) FBS and 1x Penicillin-Streptomycin (MP Biomedicals) at 37 °C with 5% (v/v) CO_2_. The cells were grown to 40% confluence and transiently transfected with the respective plasmid DNA using PEI “MAX” (MW 40,000) (Polysciences Inc, catalog no.: 24765-2) as per a protocol reported previously by us (25). For co-transfection studies to detect LPL activity, plasmid DNA of WT or S159A LPL were mixed with LMF1 in a 1:1 ratio (w/w) and were transiently transfected as mentioned earlier. Mock-transfected cells were transiently transfected with an empty vector to be used as a control for subsequent experiments. The HEK293T cells were harvested 48 to 60 h post transfection by scraping, washed with cold sterile DPBS (3x), resuspended in 500 µL of DPBS, and lysed by sonication. The overexpression of WT or S159A LPL, or LMF1 was confirmed by Western blot analysis. All Western blots were imaged on a Syngene Chemi-XRQ gel documentation system. The primary and secondary antibodies used in this Western blot analysis were anti-6X His tag antibody [HIS.H8] (Abcam, catalog no.: ab18184, dilution 1:1000) and goat anti-mouse IgG H&L (Abcam, catalog no.: ab6789, dilution 1:10000), respectively.

## RESULTS

### Profiling CE hydrolase activity in mouse tissues and blood

We recently developed a robust LC-MS based method to profile cholesterol and CE species across multiple mouse tissues and whole blood (2). Our analysis revealed that unsaturated CE species were the predominant forms present in all tissues and blood samples (2). Based on this observation, we hypothesized that tissue-associated CE hydrolases would preferentially hydrolyze unsaturated CE species. To test this, we established an *in vitro* substrate hydrolysis assay using C18:1 CE, the most abundant CE species identified across tissues, as the substrate. C18:1 CE was incorporated into PC-containing vesicles and incubated with lysates prepared from different mouse tissues.

Among all tissues examined, kidney lysates exhibited the highest specific CE hydrolase activity, followed by lung and spleen lysates (**Figure 2A**). Brain, heart, and liver tissues displayed relatively lower specific CE hydrolase activities, whereas skeletal muscle showed negligible activity (**Figure 2A**). Notably, despite having one of the highest abundance and diversities of CEs, the liver exhibited comparatively low specific CE hydrolase activity relative to other metabolically active tissues like the kidney, suggesting alternative mechanisms of utilizing CEs (**Figure 2A**).

**Figure 2.**
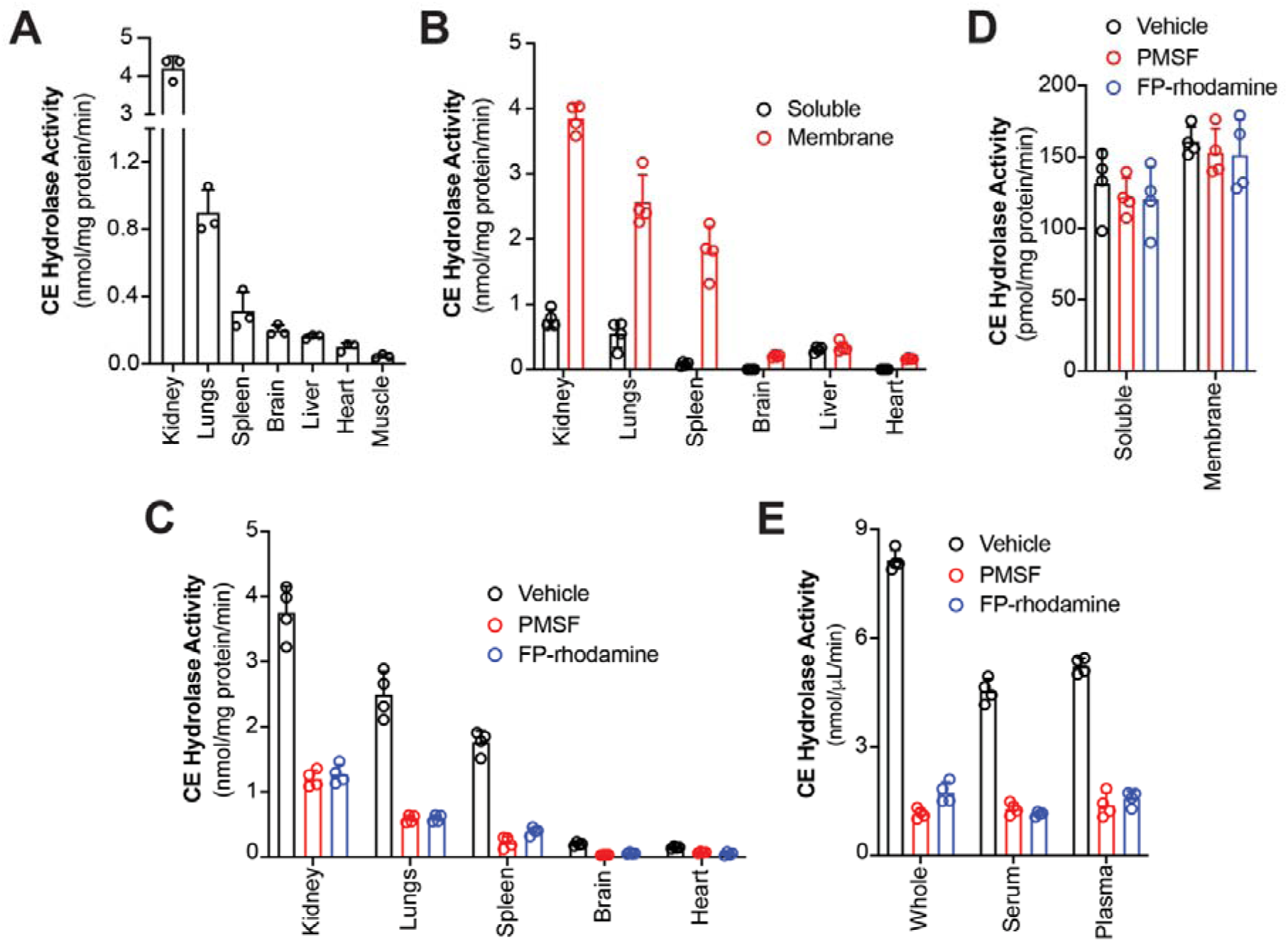
CE hydrolase activity in different mouse tissues and blood. The specific CE hydrolase activity measured in: (**A**) various mouse tissue lysates, (**B**) soluble and membrane proteomic fractions of different mouse tissues, (**C**) membrane proteomic fraction of different tissues after treatment with PMSF and FP-rhodamine, (**D**) liver soluble and membrane proteomic fractions after treatment with PMSF and FP-rhodamine, (**E**) mouse blood samples (whole blood, serum and plasma) after treatment with PMSF and FP-rhodamine. For **C, D** and **E**: PMSF treatment (2 mM, 1 h, 37 °C) and FP-rhodamine treatment (20 µM, 45 min, 37°C). All bar data is represented as mean ± standard deviation from three (for **A**) or four (for **B-E**) biological replicates per experimental group.

To further pinpoint the tissue-resident CE hydrolase activity, tissue lysates were fractionated into soluble and membrane-associated proteomes using established protocols (25). Subsequent substrate hydrolysis assays on these tissue proteomic fractions demonstrated that the CE hydrolysis activity was predominantly enriched within the membrane fractions of all tissues examined (**Figure 2B**). Interestingly, the liver represented a notable exception, where both soluble and membrane fractions exhibited comparable CE hydrolase activity levels (**Figure 2B**).

In mammals, most lipases acting on neutral lipids belong to the metabolic serine hydrolase (mSH) superfamily (29), and we wanted to assess if our lipase of interest belonged to this enzyme class. We therefore hypothesized that if the observed CE hydrolysis activity was mediated by lipase(s) from the mSH family, inhibition of this enzyme class by a broad spectrum inhibitor should reduce its activity (29). To test this, tissue membrane fractions were independently treated with two broad spectrum active-site directed irreversible inhibitors of the mSH family (29): (i) phenylmethylsulfonyl fluoride (PMSF; 2 mM, 1 h, 37°C), and (ii) fluorophosphonate-rhodamine (FP-rhodamine; 20 µM, 45 min, 37°C), prior to performing CE hydrolysis assays.

Treatment with either PMSF or FP-rhodamine resulted in greater than three-fold reductions in CE hydrolase activity across all tissue membrane fractions (except the liver), indicating that the observed activity is largely mediated by one or more mSH enzymes (**Figure 2C**). In contrast, liver CE hydrolysis activity was relatively resistant to inhibition, suggesting that the measured activity in this tissue may instead arise from one or more acyl-transferases (**Figure 2D**). This may reflect compensatory mechanisms that facilitate cholesterol to CE interconversion in the liver, potentially driven by the high ACAT activity characteristic of this tissue (30, 31). Collectively, these findings indicate that CE hydrolase activity in most mouse tissues (except the liver) is mediated predominantly by one (or more) membrane-associated mSH enzyme(s).

We next extended these analyses to mouse whole blood, as well as serum and plasma fractions. The specific CE hydrolase activity in all blood fractions, namely whole blood, serum and plasma was fairly high (**Figure 2E**). Similar to tissue-derived activities, the CE hydrolase activity in all the blood fractions were sensitive to inhibition by both PMSF and FP-rhodamine, further supporting the involvement of mSH enzymes regulating this activity in the blood (and perhaps systemically in circulation).

### Inhibitor screening of tissue membrane lysates

To identify membrane-associated mSHs that function as CE hydrolases in mouse tissues, we employed a pharmacological screening strategy using three potent active-site directed inhibitors (**Figure S1**). These included JW480, a covalent selective inhibitor of NCEH1/KIAA1363, a putative CE hydrolase (Sigma, catalog no.: SML0972-5MG) (32); Lalistat-2, a reversible inhibitor of the acidic CE hydrolase LAL, and the putative neutral CE hydrolase LIPE/HSL (Sigma, catalog no.: SML2053-5MG) (33); and tetrahydrolipstatin (THL; Orlistat), a broad-spectrum lipase inhibitor of the mSH family (Sigma, catalog no.: O4139-25MG) (25). This inhibitor panel was selected to systematically narrow down the enzyme responsible for neutral CE hydrolase activity in light of some conflicting evidence available in the literature.

Membrane proteomes isolated from kidney, lung, heart, and spleen tissues, organs that exhibited the highest specific CE hydrolase activity in our earlier profiling experiments (**Figure 2A**), were individually treated with each inhibitor (50 µM, 30 min) and subsequently assayed for CE hydrolase activity. An inhibitor was considered a “hit” if it reduced CE hydrolase activity by ≥ 50% relative to vehicle-treated controls. Among the three inhibitors tested, only THL consistently reduced the neutral CE hydrolase activity by ≥ 50% across all tissue membrane lysates examined (**Figure 3A**), suggesting that the responsible membrane-associated mSH is a THL-sensitive enzyme.

**Figure 3.**
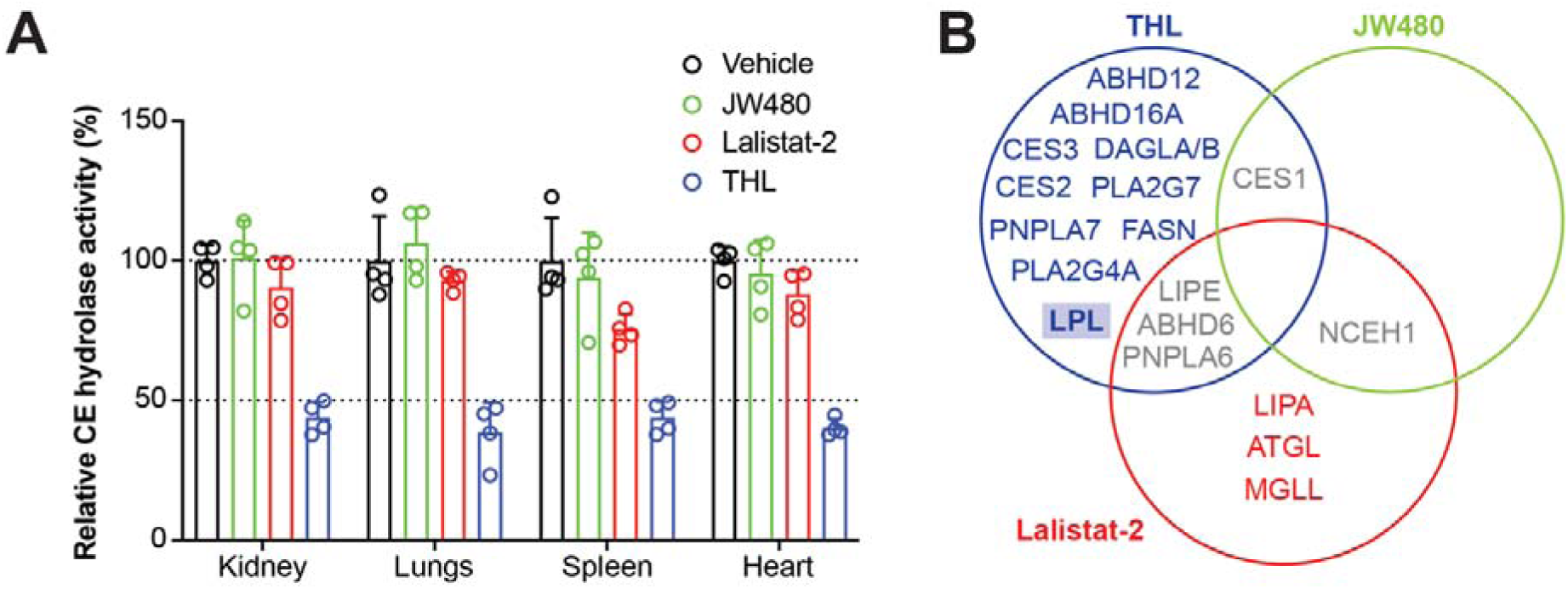
Inhibitor screening of mouse tissue membrane lysates. (**A**) Relative CE hydrolase activity of membrane proteomic fraction of mouse tissues having highest enzymatic activity, treated with vehicle or various inhibitors (JW480, Lalistat-2 or THL) (50 µM, 30 min). All bar data is represented as mean ± standard deviation from four biological replicates per experimental group. (**B**) Venn diagram analysis of the various known biological targets (lipases from the mSH family inhibited) of the three inhibitors tested in our assays.

To further refine the shortlisting of a candidate lipase, we compared the target profiles of the three inhibitors based on available literature (25). Neither JW480 nor Lalistat-2 caused a measurable reduction in CE hydrolase activity, indicating that NCEH1 and LIPE are unlikely to account for the observed activity in these tissues. Although THL is also known to inhibit LIPE, Lalistat-2 exhibits stronger inhibition of LIPE than THL (33), allowing us to reasonably exclude LIPE as the primary contributor. This filtering approach narrowed the candidate pool to THL-specific targets that do not overlap with those of JW480 or Lalistat-2 (**Figure 3B**). After excluding proteases and considering prior knowledge of the various lipases based on published mSH biology, lipoprotein lipase (LPL) emerged as the strongest candidate enzyme capable of CE hydrolysis.

### Identification of LPL as a putative CE hydrolase

We previously demonstrated that RAW264.7 cells, an immortalized murine macrophage cell line, exhibit significant accumulation of CEs accompanied by a concomitant reduction in free cholesterol following THL treatment (25 µM, 8 h), suggesting the presence of a THL-sensitive lipase with putative CE hydrolase activity (2). To directly assess this enzymatic activity, lysates from RAW264.7 cells were subjected to CE substrate hydrolysis assays following treatment with vehicle (DMSO) or THL (50 µM, 30 min). RAW264.7 lysates displayed robust CE hydrolase activity that was significantly reduced upon THL treatment, consistent with the CE and cholesterol measurements reported earlier (**Figure 4A**). We additionally screened several other cell lines for THL-sensitive CE hydrolase activity. Here, we observed that Neuro2A cells had abundant CEs and cholesterol (2), but lacked any detectable THL-sensitive CE hydrolysis activity (**Figure 4A, Figure S2**), making this cell line a suitable comparator (control) for subsequent target identification studies.

**Figure 4.**
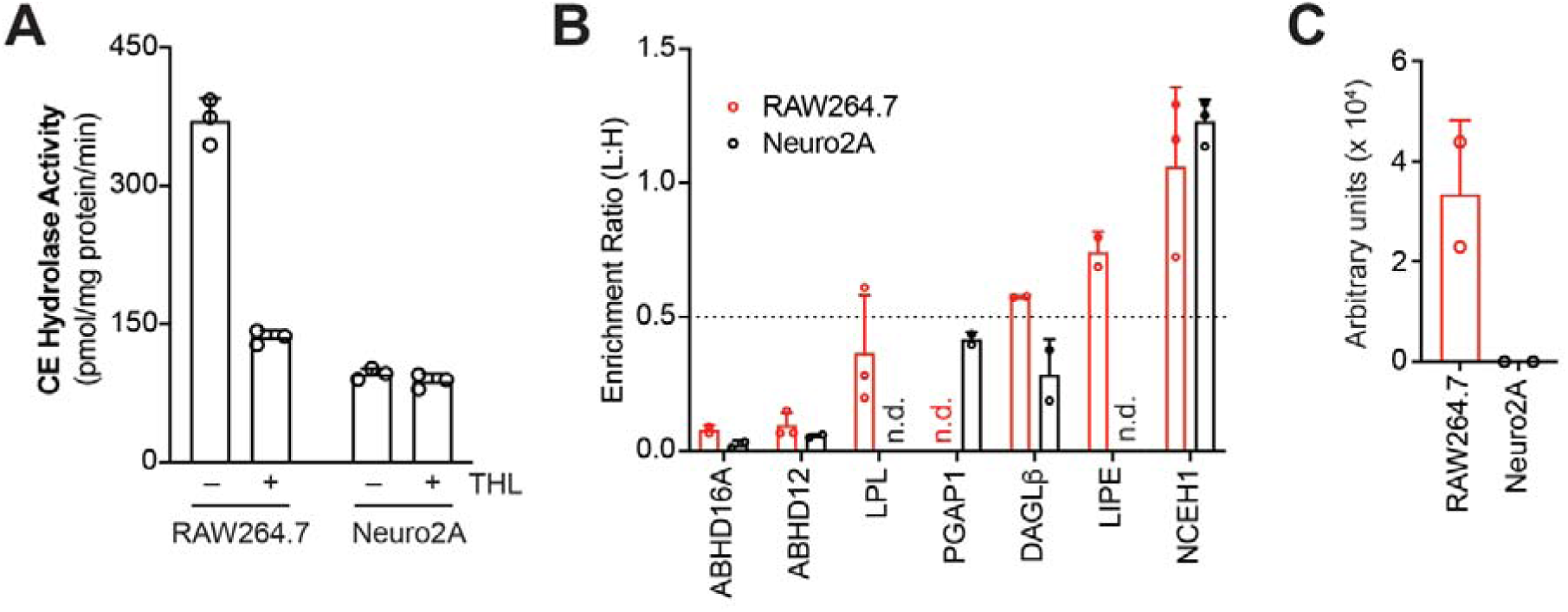
LPL as a putative CE hydrolase. (**A**) The specific CE hydrolase activity in RAW264.7 and Neuro2A cell lysates following *in vitro* treatment with vehicle (DMSO) or THL (50 µM, 30 min). (**B**) A LC-MS/MS based competitive ABPP experiment showing the enrichment ratio (light:heavy, L:H) of the mSHs inhibited by THL treatment (50 µM, 30 min) (by ≥ 50%) of RAW264.7 (red) or Neuro2A (black) cells. The horizontal dotted line denotes an enrichment ratio 0.5, and mSHs having an enrichment ratio of ≤ 0.5 were considered as hits (THL-sensitive mSHs) in this study. The putative CE hydrolases described in literature, i.e. LIPE and NCEH1, were not significantly inhibited by THL in RAW264.7 cells, and hence, not considered as a potential CE hydrolase in subsequent studies. Complete enrichment ratios of all mSHs identified in this study can be found in **Table S1**. Here, n.d. means not detected. (**C**) mRNA levels of LPL in RAW264.7 and Neuro2A cells directly taken from a publicly available large-scale gene expression databased (https://biogps.org) (38, 39). All bar data is represented as mean ± standard deviation from three (for **A, B**), and two (for **C**) biological replicates per experimental group.

Both RAW264.7 and Neuro2A cells possess a rich diversity of lipases from the mSH family (34, 35), and the differential CE hydrolase activity observed between these cell lines prompted us to identify the underlying lipases from this family using established competitive activity-based protein profiling (ABPP) methods (23). RAW264.7 and Neuro2A cells were individually treated with THL (25 µM, 8 h) or vehicle control, followed by enrichment of mSHs with FP-biotin (10 µM, 1 h) and their downstream identification using reported LC-MS/MS analysis (26). A mSH enzyme was classified as a THL target if it was identified in at least two of three biological replicates, quantified with one or more peptides per biological replicate, and exhibited an average light-to-heavy (L:H) ratio ≤ 0.5, corresponding to > 50% inhibition by THL.

Using this approach, we identified 41 and 31 mSHs in RAW264.7 and Neuro2A cells, respectively, with 17 and 7 uniquely detected in RAW264.7 and Neuro2A cells respectively (**Table S1, Figure S3**). Application of the filtering criteria identified three and four THL sensitive lipases from the mSH in RAW264.7 and Neuro2A cells respectively (**Figure 4B**). ABHD12 and ABHD16A, both known targets of THL (36, 37), were excluded because they were detected and equally inhibited in both cell lines. The only remaining lipase uniquely inhibited by THL in RAW264.7 cells was LPL. Consistent with these observations, publicly available gene expression datasets revealed strong LPL expression in RAW264.7 cells but negligible expression in Neuro2A cells (**Figure 4C**) (38, 39). Taken together, these findings strongly corroborated our inhibitor-screening data from mouse tissues, and identify LPL as a promising candidate CE hydrolase in mammalian cells and tissues. LPL is a membrane-associated homodimeric enzyme (∼110 kDa) that has been classically characterized as a triacylglycerol lipase (40–42). Although previous studies have shown that LPL can bind CEs (40–42), its role as a CE hydrolase has not been investigated to the best of our knowledge.

To further evaluate the role of LPL in CE hydrolysis, we initially tested GSK264220A (Sigma, catalog no.: SML2931-5MG) (**Figure S1**), a covalent inhibitor of endothelial lipase (LIPG) reported to exhibit off-target activity against LPL (43). Treatment of RAW264.7 cells with GSK264220A (50 µM, 4 h) did not alter CE or cholesterol levels (**Figure S4**). Similarly, cellular lysates from RAW264.7 cells treated with GSK264220A (50 µM, 30 min) did not affect the CE hydrolase activity (**Figure S4**). Competitive ABPP profiling following GSK264220A treatment (10 µM, 8 h) of RAW264.7 cells identified 39 mSHs, but none exhibited any appreciable inhibition (**Table S1, Figure S4**), suggesting that GSK264220A is a poor inhibitor of LPL under our experimental conditions, and cannot be used in cells to ascertain LPL activity.

### LPL functions as a CE hydrolase *in vitro*

To directly evaluate the role of LPL as a CE hydrolase, we transiently overexpressed mouse wild-type (WT) LPL in HEK293T cells using a previously reported transfection strategy (23, 25). Western blot analysis confirmed robust expression of WT LPL relative to mock-transfected controls (**Figure S5**). However, gel-based ABPP and CE hydrolase assays revealed that the overexpressed WT LPL was catalytically inactive (**Figure S5**). Since ABPP based assessment of LPL overexpression had not been previously reported, we examined earlier studies describing LPL expression in HEK293T cells. These reports showed that mouse LPL requires the endoplasmic reticulum chaperone lipase maturation factor 1 (LMF1) for proper folding, dimerization, and enzymatic activity, and that HEK293T cells lack endogenous LMF1 expression (44, 45). Thus, the absence of LMF1 likely accounted for the inactivity of overexpressed LPL in our initial experiments.

To restore LPL activity, we co-transfected HEK293T cells with mouse WT LPL and mouse LMF1. As a catalytic control, we generated an active-site variant of LPL in which the catalytic Ser-159 residue was mutated to alanine (S159A), and co-expressed this mutant together with LMF1. Western blot analysis demonstrated comparable expression levels of WT and S159A LPL relative to mock-transfected controls (**Figure 5A**). In parallel, cells transfected with LMF1 alone also showed robust LMF1 expression (**Figure 5A**). Gel-based ABPP analysis revealed strong enzymatic activity exclusively in cells expressing WT LPL, whereas neither the S159A variant of LPL nor LMF1 alone exhibited detectable activity (**Figure 5A**).

**Figure 5.**
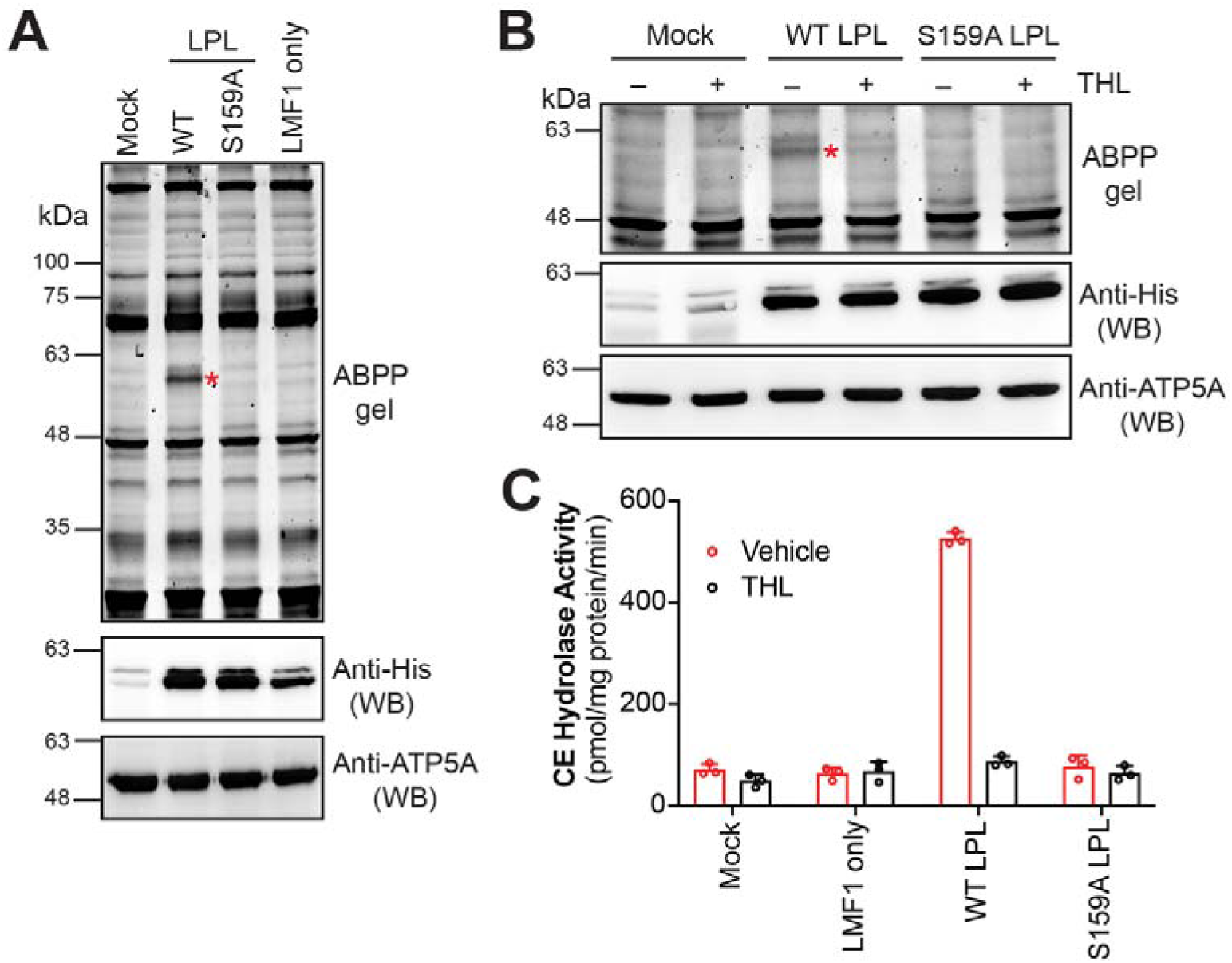
WT LPL possesses CE hydrolase activity *in vitro*. (**A**) HEK293T cell lysates transfected with mock control (empty plasmid), LMF1 only (control), WT LPL + LMF1, or S159A LPL + LMF1 were assessed using gel-based ABPP (top panel) and Western blot (WB) analysis using an anti-His antibody (to confirm overexpression of LPL variants and LMF1) or an anti-ATP5A antibody (to confirm equal loading of samples). The red asterisk corresponds to the activity (top panel) of WT LPL in the gel-based ABPP experiment. LMF1 and LPL have similar molecular weights and hence so overlapping bands in the WB experiments. This experiment was done three times with reproducible result each time. Complete images of the WB experiments can be found in the ***Supporting Information***. (**B**) HEK293T cell lysates transfected with mock control (empty plasmid), WT LPL + LMF1, or S159A LPL + LMF1 were assessed using gel-based ABPP (top panel) after treatment with THL (50 µM, 30 min at 37°C). To confirm intactness of LPL and equal loading post-treatment, Western blot (WB) analysis using an anti-His antibody (to confirm overexpression of LPL variants and LMF1) or an anti-ATP5A antibody (to confirm equal loading of samples) were performed. The red asterisk corresponds to the activity (top panel) of WT LPL in the gel-based ABPP experiment. This experiment was done three times with reproducible result each time. Complete images of the ABPP and WB experiments can be found in the ***Supporting Information***. (**C**) The specific CE hydrolase activity of HEK293T cell lysates transfected with mock control (empty plasmid), LMF1 only (control), WT LPL + LMF1, or S159A LPL + LMF1 after treatment with vehicle (DMSO) or THL (50 µM, 30 min at 37°C). All bar data is represented as mean ± standard deviation from three biological replicates per experimental group.

To pharmacologically validate LPL activity, lysates from HEK293T cells expressing WT LPL + LMF1, S159A LPL + LMF1, or mock controls were treated with THL (50 µM, 30 min). Gel-based ABPP assays demonstrated that THL substantially inhibited WT LPL activity, while no effect was observed in the catalytically inactive S159A variant (**Figure 5B**). Western blot analysis confirmed that THL treatment altered only LPL activity and did not affect protein expression levels of either WT or S159A LPL variants (**Figure 5B**).

We next performed CE hydrolase assays using lysates from HEK293T cells expressing WT or S159A LPL together with LMF1. Relative to mock-transfected controls, only WT LPL exhibited robust CE hydrolysis activity, whereas the S159A mutant showed no detectable enzymatic activity (**Figure 5C**). Cells expressing LMF1 alone also lacked CE hydrolase activity and were comparable to mock controls (**Figure 5C**). Importantly, treatment of WT LPL expressing lysates with THL (50 µM, 30 min at 37°C) resulted in significant reduction in CE hydrolase activity, while no significant changes were observed in S159A LPL or LMF1-only controls (**Figure 5C**).

The requirement of LMF1 for LPL activation highlights an additional regulatory layer governing LPL function. Analysis of publicly available gene expression (38, 39) and proteomics datasets (35, 46) revealed that LPL and LMF1 are not uniformly expressed across tissues (**Figure S6**), which may contribute to the tissue-specific variability in CE hydrolase activity observed *in vivo*. Collectively, these biochemical and pharmacological studies demonstrate that LPL indeed functions as a robust CE hydrolase *in vitro* and that its activity is sensitive to inhibition by THL. These findings strongly support our cellular and inhibitor-screening data identifying LPL as a mammalian CE hydrolase.

## DISCUSSION

Cholesterol is an essential lipid in mammals, where it regulates a broad range of physiological processes including metabolism, membrane organization, and cellular signaling (1, 47). Excess cellular cholesterol is converted into CEs, which serve as inert storage and transport forms of cholesterol (2). Cellular cholesterol homeostasis therefore depends on a tightly regulated balance between cholesterol esterification and CE hydrolysis. Although the enzymes responsible for cholesterol esterification, namely ACAT and LCAT, have been extensively characterized, comparatively little is known about the enzymes that hydrolyze CEs back to free cholesterol. Several candidate CE hydrolases have been reported; however, among these, only LAL, an acidic CE hydrolase, has been functionally validated (16, 17). In contrast, the CE hydrolase activity of neutral lipases, including LIPE and NCEH1, remain controversial (18–22), suggesting that additional enzymes contributing to neutral CE hydrolysis likely remain unidentified.

To address this gap, we systematically profiled CE hydrolase activity across mouse tissues and blood. CE hydrolase activity was detected in nearly all tissues examined (with the exception of muscle), and in blood samples. Tissue fractionation and broad-spectrum inhibitor treatments further demonstrated that the predominant CE hydrolase activity was coming from one (or more) membrane-bound mSH enzyme(s). Using an additional inhibitor-based screening strategy, we found that the putative CE hydrolase activity was highly sensitive to THL. Combining substrate hydrolysis assays with competitive ABPP profiling in RAW264.7 and Neuro2A cells enabled us to identify a THL sensitive target selectively enriched in RAW264.7 cells. This analysis ultimately implicated LPL, a homodimeric membrane-associated enzyme, as a candidate CE hydrolase (40–42). Consistent with this conclusion, *in vitro* biochemical assays demonstrated that WT LPL, but not the catalytically inactive S159A mutant, robustly hydrolyzed CEs. Importantly, LPL activity required the presence of LMF1 (44, 45), as WT LPL expressed in the absence of LMF1 was catalytically inactive. Collectively, these findings establish LPL as a functional CE hydrolase and suggest that tissue-specific regulation of the LPL-LMF1 axis may contribute significantly to CE metabolism *in vivo*.

Our findings also reveal that CE metabolism is likely mediated by distinct enzymatic mechanisms in different tissues. In the liver, CE hydrolase activity was detected in both soluble and membrane fractions; however, this activity was largely insensitive to FP-probe treatment. This observation suggests that the hepatic CE turnover observed in our assays may not primarily involve mSHs and could instead reflect the action of acyltransferases or related enzymes. Given the high expression of ACAT1 and the additional presence of ACAT2 in liver, it is plausible that enhanced esterification and lipoprotein-mediated CE efflux substantially contribute to the observed substrate turnover (3, 4, 30, 31). Similarly, although mouse blood, plasma, and serum exhibited strong CE hydrolysis activity, FP-probes did not completely abolish this activity, indicating that non-mSH enzymatic mechanisms may also contribute to the observed CE hydrolase activity. These findings raise the intriguing possibility that acyltransferase-dependent remodeling pathways cooperate with LPL-mediated hydrolysis to regulate CE turnover in circulation (48).

Interestingly, spleen tissue exhibited strong CE hydrolase activity that was sensitive to both FP-probes and THL, despite lacking detectable expression of LPL and LMF1 (**Figure S6**) (38, 39). This finding suggests the existence of additional, as of yet unidentified, membrane-associated mSH(s) capable of mediating CE hydrolysis in immune tissues. Identifying these enzymes will be important for understanding tissue-specific CE metabolism and may reveal additional regulatory pathways involved in cholesterol homeostasis.

Our study also raises several broader implications regarding the physiological functions of LPL. LPL is well established as a triglyceride lipase and has been implicated in the uptake of CEs through cholesteryl ester transfer protein (CETP)-dependent mechanisms (49); however, a direct role for LPL in CE hydrolysis had not previously been demonstrated. The identification of LPL as a CE hydrolase suggests that it may contribute directly to CE turnover during lipoprotein remodeling, including HDL to LDL conversion. Because LPL activity is tightly regulated in a tissue-specific manner through multiple transcriptional and post-translational mechanisms, CE hydrolysis is also likely to be dynamically regulated across tissues and physiological states. This possibility warrants further investigation.

Notably, LPL knockout mice exhibit severe dyslipidaemia and die within 1 to 2 days after birth, underscoring the central importance of LPL in systemic lipid metabolism (50). Conversely, LPL overexpression reduces circulating triglyceride and cholesterol levels and markedly attenuates atherogenesis (51). These observations raise the possibility that the protective effects of LPL may arise not only from its established triglyceride lipase activity but also from its newly identified role in CE hydrolysis. More broadly, our findings help address the longstanding question surrounding the identity of neutral CE hydrolases and provide a framework for investigating CE metabolism in both physiological and pathological settings. In particular, understanding how LPL and related enzymes regulate oxidized CEs may offer new insight into lipid dysregulation during oxidative stress and diseases such as atherosclerosis (52, 53).

## CONCLUSION

In summary, our study identifies LPL as a previously unrecognized neutral CE hydrolase in mammalian systems. Through tissue-wide activity profiling, pharmacological inhibition, competitive chemoproteomics, and biochemical validation, we demonstrate that LPL hydrolyzes CEs in a catalytic and THL-sensitive manner. Importantly, this activity requires LMF1, highlighting a regulatory axis that may govern tissue-specific CE turnover. These findings address a longstanding gap in cholesterol biology by providing molecular evidence for a physiologically relevant neutral CE hydrolase distinct from previously proposed candidates like LIPE and NCEH1. Our work further expands the known functional repertoire of LPL beyond triglyceride metabolism and suggests broader roles in lipoprotein remodeling and cholesterol homeostasis. The discovery of a LPL mediated CE hydrolysis opens new avenues for investigating lipid metabolism in physiological and pathological contexts, including atherosclerosis, metabolic disorders, and oxidative lipid signaling.

## ACCESSION CODES

The UniProt ID of mouse LPL and LMF1 are P11152 and Q3U3R4 respectively. Additionally, accession IDs of all the mSHs identified in this study can be found in Supplementary Table S1.

## SUPPORTING INFORMATION

The supporting information associated with this manuscript includes: (1) Supplementary Figures (PDF), and (2) Supplementary Table S1 (XLSX).

## DATA AVAILABILITY

All data that supports the findings of this study are available in this paper and its Supplementary Information or are available with the corresponding author (S.S.K.) upon reasonable request. All the raw proteomics data have been deposited with the ProteomeXchange Consortium via the PRIDE partner repository with the identifiers: PXD077540.

## AUTHOR CONTRIBUTIONS

A.C. and S.S.K. conceived the project, analyzed the data, and co-wrote the manuscript. A.C. performed all the experiments. S.S.K. acquired funding and supervised the project.

## ACKNOWLEDGEMENTS

We thank Saddam Shekh for maintenance of the biological mass spectrometry facility at IISER Pune, and for technical assistance. The staff at the NFGFHD at IISER Pune are thanked for helping with the mice experiments described in this study. Members of the S.S.K. are thanked for providing critical comments and inputs throughout the course of this study. We acknowledge generous financial support from the Anusudhan National Research Foundation (ANRF), Government of India (Grants: SwarnaJayanti Fellowship SB/SJF/2021-22/01 to S.S.K.). A.C. is supported by graduate student fellowship from IISER Pune.

