## Supplementary Figures for "The Hunt for Cholesteryl Ester Hydrolases: Identification of Lipoprotein Lipase as a Cholesterylesterase"

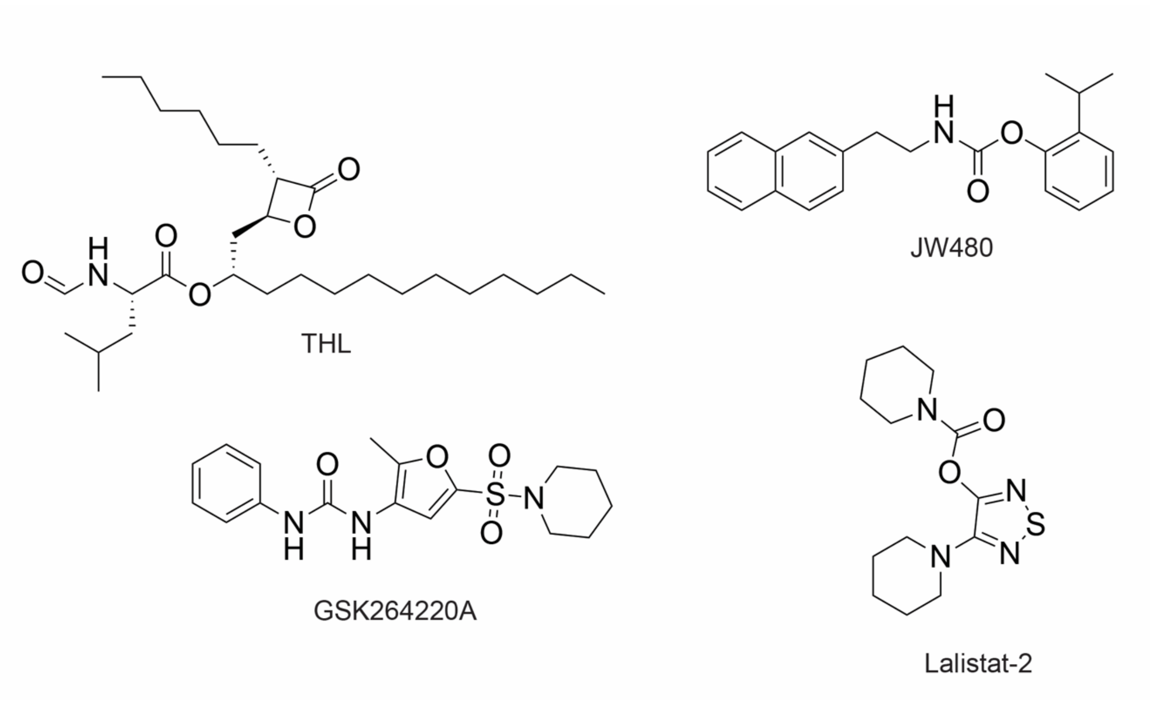


**Figure S1**. Structures of inhibitors used in this study

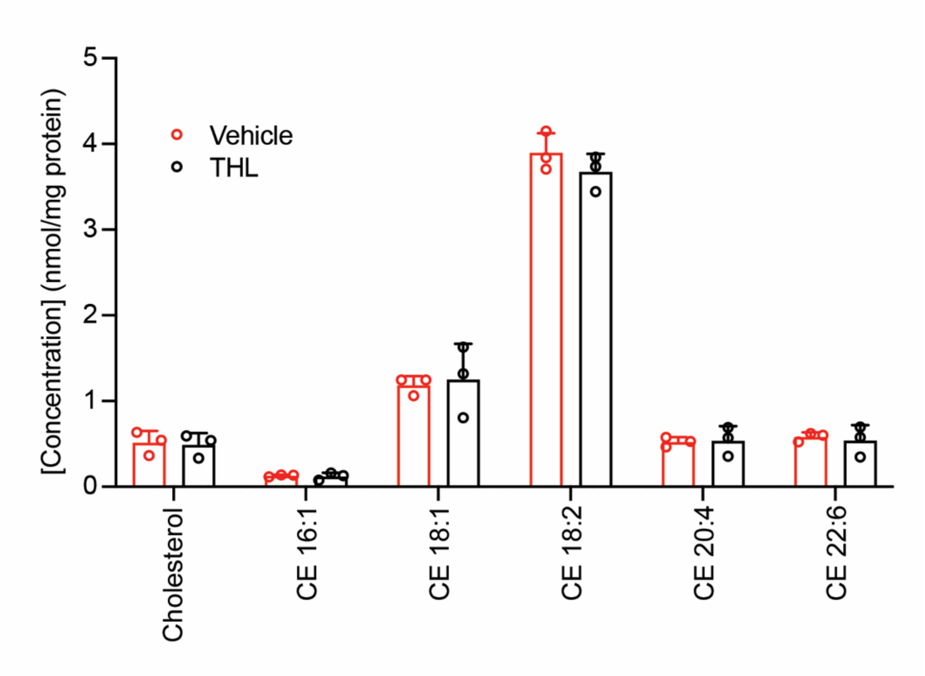


**Figure S2**. The cellular concentration of cholesterol and CEs in Neuro2A cells after treatment with THL (25 μM, 8 h), showing no changes in the levels of these lipids. All bar data is represented as mean ± standard deviation from three biological replicates per experimental group.

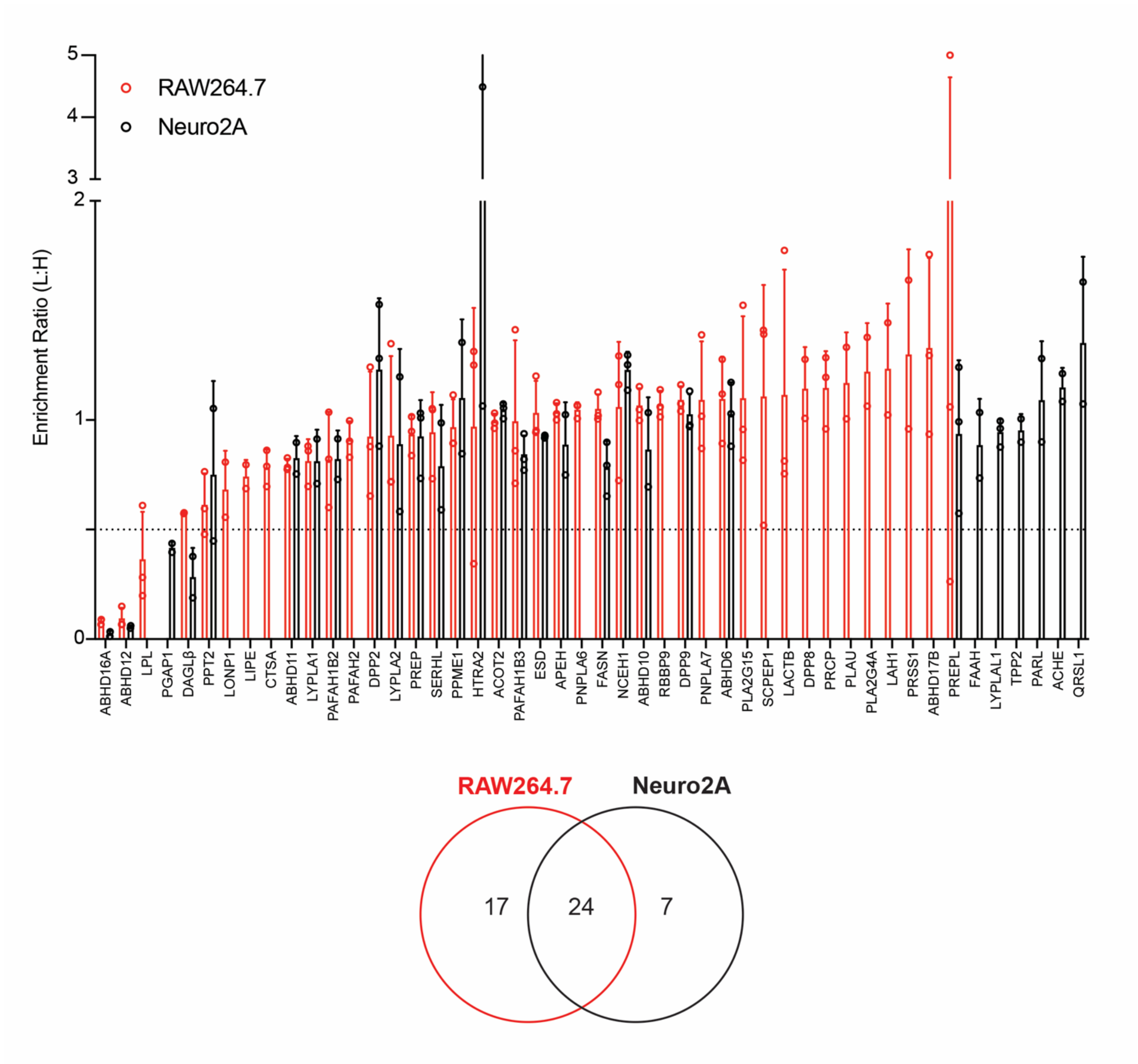


**Figure S3.** _Top panel_: Complete LC-MS/MS-based ABPP analysis for the data shown in Figure 4B. _Bottom panel_: Venn diagram showing the overall of mSHs identified in this experiment between the two cell lines tested.

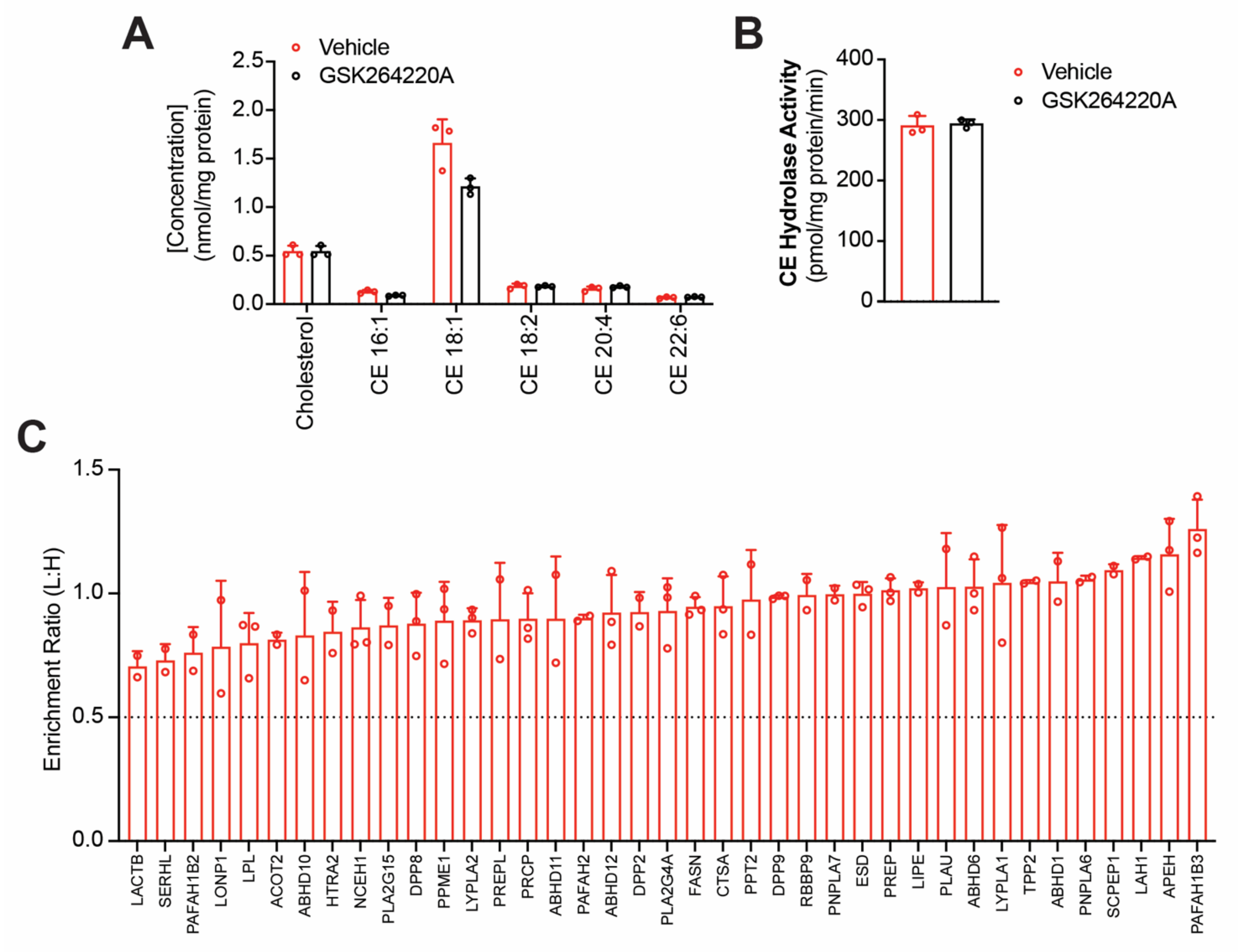


**Figure S4. GSK264220A does not inhibit LPL in RAW264.7 cells.** (**A**) The cellular concentration of cholesterol and CEs in RAW264.7 cells after treatment with GSK264220A (50 µM, 4 h), showing no changes in the levels of these lipids. (**B**) The specific CE hydrolase activity in RAW264.7 cell lysates following *in vitro* treatment with vehicle (DMSO) or GSK264220A (50 µM, 30 min), showing no change in CE hydrolase activity. (C) A LC-MS/MS based competitive ABPP experiment showing the enrichment ratio (light:heavy, L:H) of the mSHs inhibited by GSK264220A treatment (50 µM, 30 min) of RAW264.7 cells. The horizontal dotted line denotes an enrichment ratio 0.5, and mSHs having an enrichment ratio of ≤ 0.5 were considered as hits (mSHs inhibited by GSK264220A) in this study. There were no hits identified from this study. Complete enrichment ratios of all mSHs identified in this study can be found in **Table S1**. For (**A, B, C**): All bar data is represented as mean ± standard deviation from three biological replicates per experimental group.

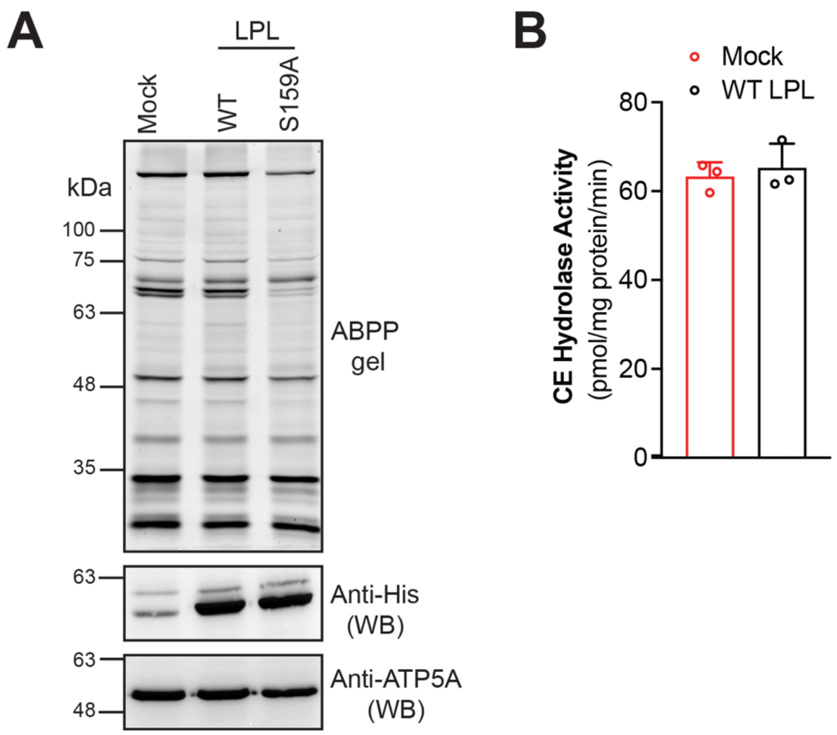


**Figure S5. WT LPL does not possesses CE hydrolase activity in the absence of LMF1.** (**A**) HEK293T cell lysates transfected with mock control (empty plasmid), WT LPL, or S159A LPL were assessed using gel-based ABPP (top panel) and Western blot (WB) analysis using an anti-His antibody (to confirm overexpression of LPL variants) or an anti-ATP5A antibody (to confirm equal loading of samples). WT LPL does not show any activity in this gel-based ABPP assay. This experiment was done three times with reproducible result each time. Complete images of the WB experiments can be found at the end of the ***Supporting Information***. (**B**) The specific CE hydrolase activity of HEK293T cell lysates transfected with mock control (empty plasmid), or WT LPL, showing no difference in activity upon LPL overexpression. All bar data is represented as mean ± standard deviation from three biological replicates per experimental group.

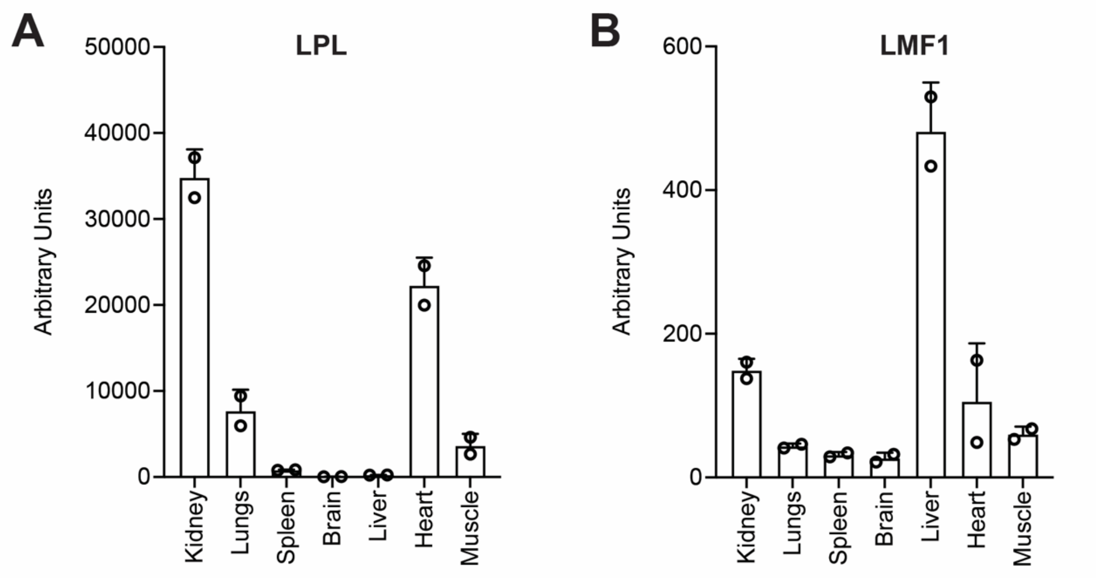


**Figure S6.** mRNA levels of LPL and LMF1 in various mouse tissues directly taken from a publicly available large-scale gene expression databased (https://biogps.org). All bar data is represented as mean ± standard deviation from two biological replicates per experimental group.

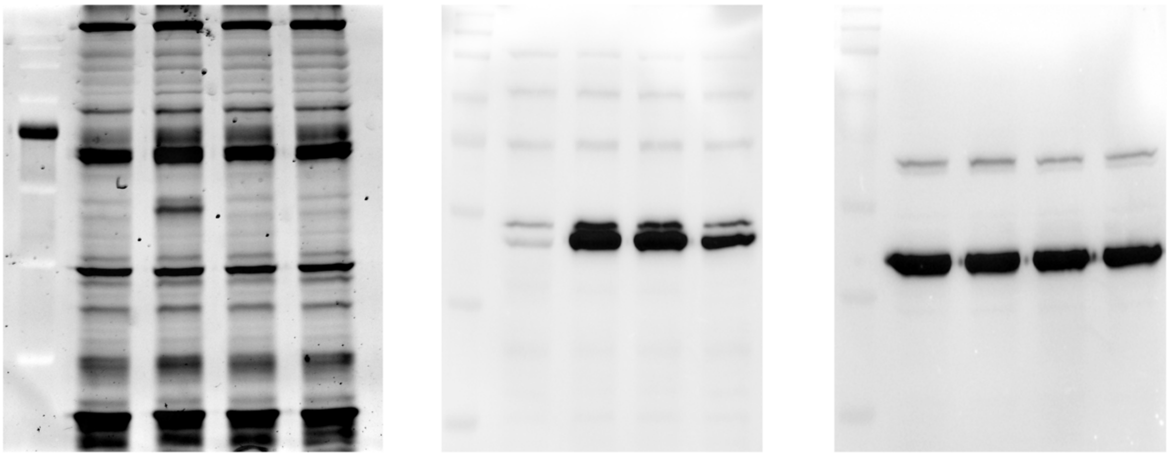

ABPP anti-His blot anti-ATP5A blot

Complete gel and blot images for Figure 5A.

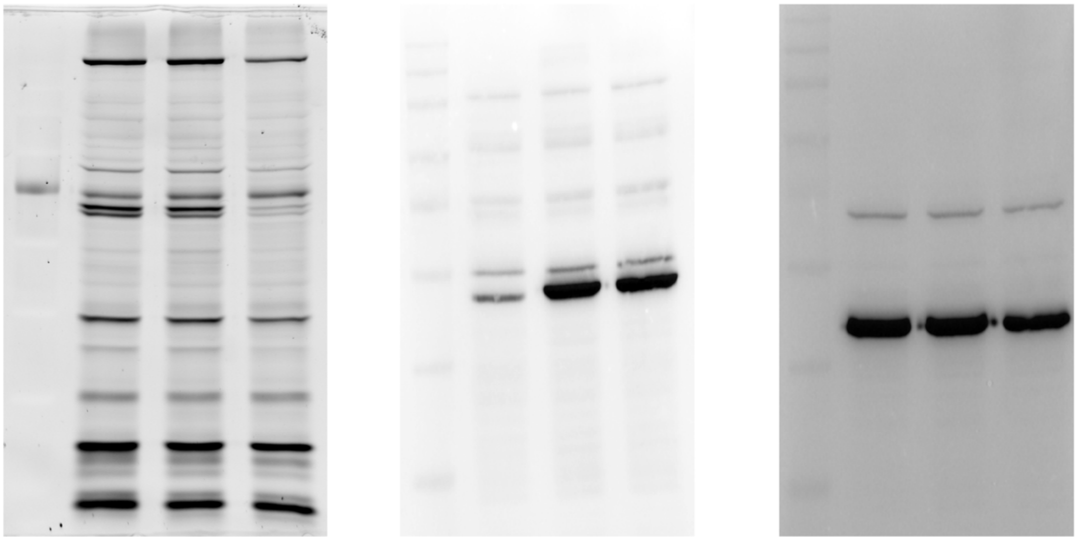

ABPP anti-His blot anti-ATP5A blot

Complete gel and blot images for Figure S5A.

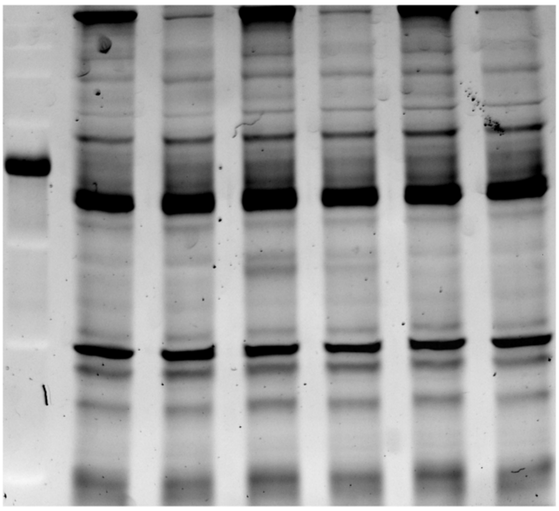

ABPP

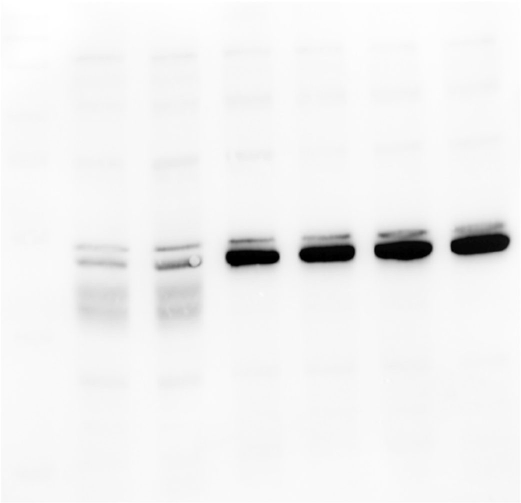

anti-His blot

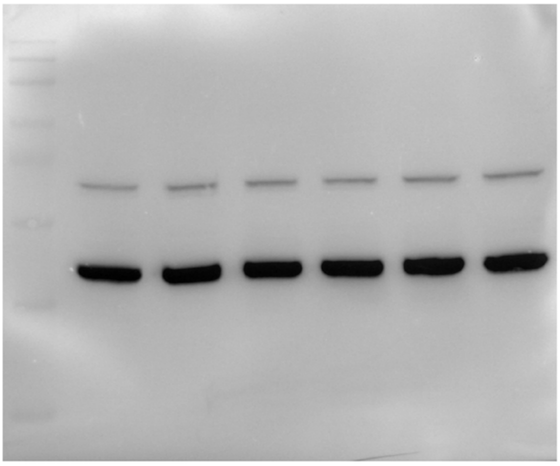

anti-ATP5A blot

Complete gel and blot images for Figure 5B.
